# CenIR, an essential BlaIR-family regulatory system in *C. difficile*

**DOI:** 10.64898/2026.04.08.717257

**Authors:** Micaila P. Kurtz, Ute Müh, David S. Weiss, Craig D. Ellermeier

## Abstract

The CenIR regulatory system of *Clostridioides difficile* comprises a predicted transcriptional repressor, CenI, and a predicted membrane metalloprotease, CenR. The physiological role of CenIR and activating signal(s) are not known. CenIR belongs to the BlaIR family of regulators that mediate resistance to β-lactam antibiotics. In canonical BlaIR systems, binding of a β-lactam to the extracellular transpeptidase domain of BlaR triggers proteolysis of BlaI and thus induction of a closely linked β-lactamase gene. However, CenR lacks a β-lactam-binding domain and transposon mutagenesis indicated CenI is essential for viability even when β-lactams are not present. Here we confirmed essentiality of CenIR and determined its regulon contains ∼12 genes, including an exported protein of unknown function (CDR_0474) that is induced about 500-fold and a peptidoglycan hydrolase (Cwp6) that is induced about 7-fold when cells are depleted of CenIR. There are no essential genes or β-lactamases in the regulon. Phenotypic characterization of CenIR-depletion strains revealed slower growth, mild elongation and cell lysis. Deletion of cdr_0474 corrected all three defects, while deletion of cwp6 only rescued the lysis phenotype. It was possible to delete *cenIR* in either a Δ*cdr_0474* or Δ*cwp6* background. We propose that CenIR is essential because its absence leads to lysis due to Cwp6 overproduction. Bioinformatic analyses revealed the predicted extracellular sensing domains in annotated “BlaR” proteins are diverse. Thus, BlaIR systems are not dedicated to defense against β-lactams but probably enable bacteria to adapt to a variety of environmental stimuli.

**Importance:** Many of the regulatory systems for controlling cell envelope biogenesis and stress responses have yet to be studied. Here we characterize a *Clostridioides difficile* BlaIR-like regulatory system that we have named CenIR for cell envelope. Unlike canonical BlaIR systems, which bind β-lactams and induce a β-lactamase, CenIR lacks a β-lactam binding domain and is essential for viability even in the absence of antibiotics. We identified the genes in the regulon and found that CenIR is essential because its absence leads to overproduction of the Cwp6 peptidoglycan hydrolase. We also show that most annotated BlaIR-like systems lack a β-lactam-binding domain, from which we infer that these systems have much broader physiological roles than generally appreciated.

## Introduction

*Clostridioides difficile* is a Gram-positive, spore-forming gastrointestinal pathogen that causes symptoms ranging from mild diarrhea to lethal pseudomembraneous colitis (1). C. difficile resides in a complex environment and encodes a large number of regulatory circuits, including roughly 50 two-component signaling systems (2). There are also seven genes annotated as BlaR-like, four of which are adjacent to a BlaI homolog. BlaIR systems are widespread in Gram-positive bacteria and have been especially well-studied in *Bacillus lichenoformis* and *Staphylococcus aureus*, where the system is known as Mec (3). In these systems, BlaI is a transcriptional repressor and BlaR is a membrane metalloprotease with an extracellular penicillin-binding transpeptidase (TP) domain (PF00905). Binding of a β-lactam to the TP domain turns on BlaR’s latent protease activity, resulting in cleavage of BlaI and induction of the target β-lactamase. Only one of *C. difficile’s* BlaIR-like systems has been characterized; it responds to β-lactams like a canonical BlaIR system (4).

BlaIR systems are not essential in any organism where they have been studied, nor would essentiality be expected in the absence of β-lactams. We were therefore intrigued when a gene pair annotated as “*blaIR-like*” was identified as essential for vegetative growth in a large-scale transposon mutagenesis of *C. difficile* strain R20291 (5). The genes in question are the blaI-like cdr_0472 and the blaR-like cdr_0473. (Note that locus names in this report have been shortened from *cdr20291_xxxx* to *cdr_xxxx*). The BlaR-like protein lacks a TP domain, suggesting it does not sense β-lactam antibiotics but contains the membrane metalloprotease domain that likely cleaves BlaI in response to a signal. Moreover, essentiality under standard laboratory growth conditions argues for a role other than (or at least beyond) mounting a stress response.

Here, we report an initial characterization of this gene pair, which we have named cenI (*cdr_0472*) and cenR (*cdr_0473*) because cells lacking CenIR induce expression of multiple <u>c</u>ell <u>en</u>velope genes, elongate and lyse.

## Results

The genes *cenI* and *cenR* are predicted to be in a two-gene operon with one transcriptional start site (as described for strain 630 (6); Fig. 1A). Like canonical BlaI proteins, CenI has a predicted helix-turn-helix DNA-binding motif (3). CenR, like canonical BlaR proteins, is predicted to have a cytoplasmic N-terminal Peptidase_M56 domain that is anchored in the membrane through 4 transmembrane helices. However, the C-terminal TP domain characteristic of canonical BlaR proteins is replaced in CenR with a shorter domain (by ∼100 amino acids) that is predicted to be largely disordered (Fig. 1). Thus, the predicted structures of CenI and CenR suggest CenI binds as a dimer to target genes to repress their expression. Interaction of something other than a β-lactam with CenR’s extracellular sensing domain triggers proteolysis of CenI and gene induction.

**FIG 1.**
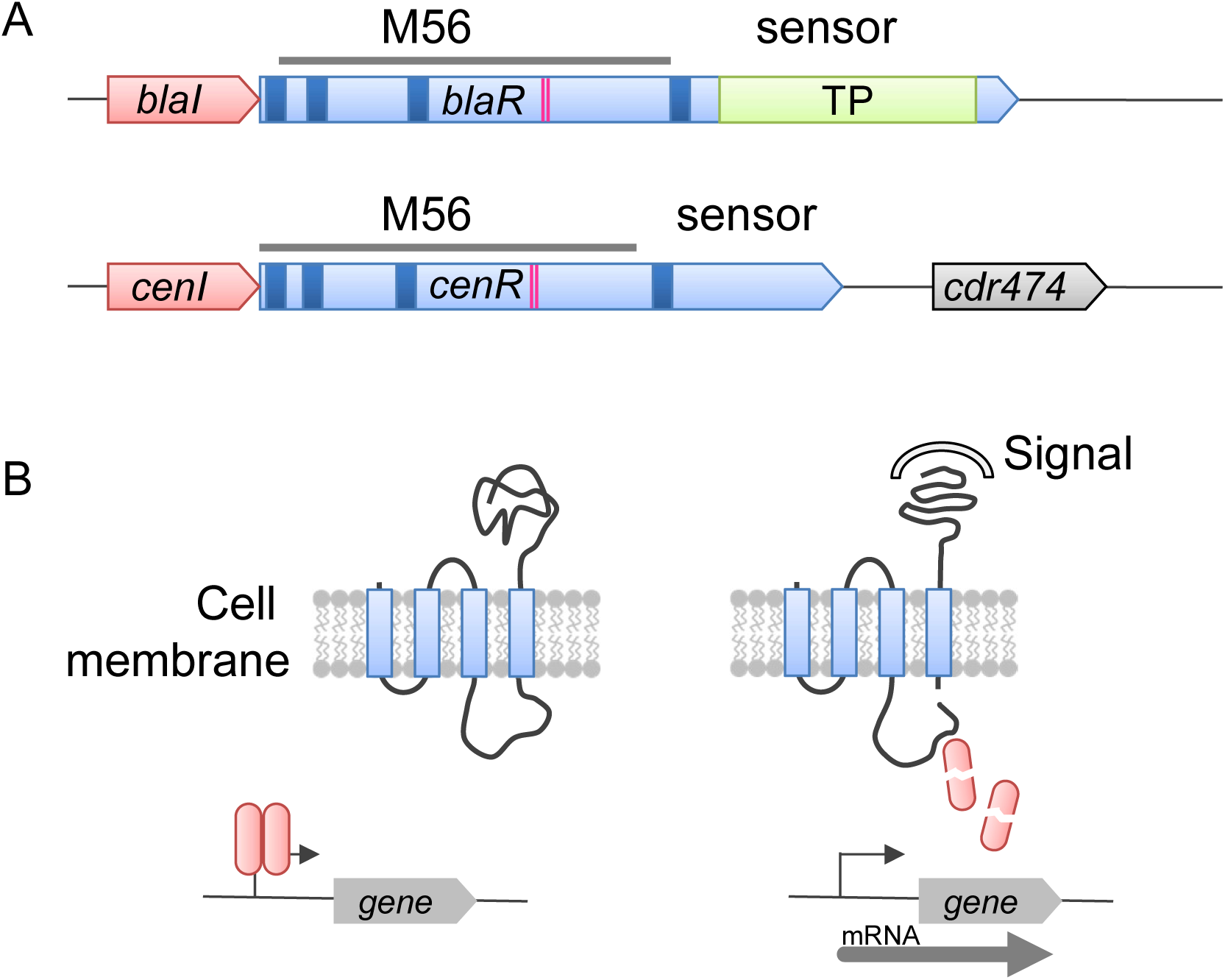
Model of CenIR operon and regulation. (A) Operon structure of *cenIR* and comparison to *blaIR*. Genes *cenI* and *blaI* (red) encode a transcriptional repressor that binds DNA as a dimer. Genes *cenR* and *blaR* (blue) encode a sensor protease with an N-terminal M56 peptidase domain that includes 4 transmembrane helices (dark blue) and zinc-binding histidines (magenta). The C-terminal end of BlaR has a transpeptidase domain (TP, green) that binds β-lactams. In contrast, the C-terminal part of CenR is about 100 amino acids shorter and not homologous to any known domain. AlphaFold modeling predicts the C-terminus of CenR to be largely disordered. (B) Model of regulation based on analogy to BlaR. CenR’s disordered domain binds to an unknown signal, inducing a conformational change that activates autoproteolysis near the 4^th^ TM, followed by cleavage of CenI, and induction of the regulon.

### CenIR-depletion slows growth, increases cell length and promotes lysis

Initially, we attempted to delete *cenIR* using CRISPR mutagenesis but were unsuccessful. The same CRISPR editing plasmid was later used to delete *cenIR* in a different strain background (described below), suggesting the inability to delete *cenIR* in wild type is due to biological rather than technical reasons. Instead, we constructed a depletion strain by using CRISPR to insert a P_tet_ promoter in front of the *cenIR* operon. The P*_tet_*::*cenIR* depletion strain grew well in the presence of the inducer anhydrotetracycline (aTet), but growth slowed upon transfer to media lacking aTet (Fig. 2A). Depleted cells were longer than WT and lysed faster in an in vitro assay in which cells are suspended in phosphate buffer containing 0.01% Triton X-100 (Fig. 2B,C). Note that the rate of lysis is dependent on the growth phase when cells are harvested (Fig. S1); care was therefore taken to harvest cells for the lysis assay at the same OD_600_ of about 0.6-0.8 throughout this report. All phenotypic changes were restored to wild-type when cenI was expressed from a plasmid under P*_xyl_* control (Fig. 2). Elongation and lysis were also observed in time-lapse microscopy of live cells of the P*_tet_*::*cenIR* strain growing under depletion conditions on an agarose pad (Fig. 2D, Videos S1-4). Curiously, however, in spot titer assays in the absence of aTet, the depletion strain grew out to the same dilution as a WT control (Fig. S2A). We had expected a more profound viability defect in view of our inability to delete cenIR.

**FIG 2.**
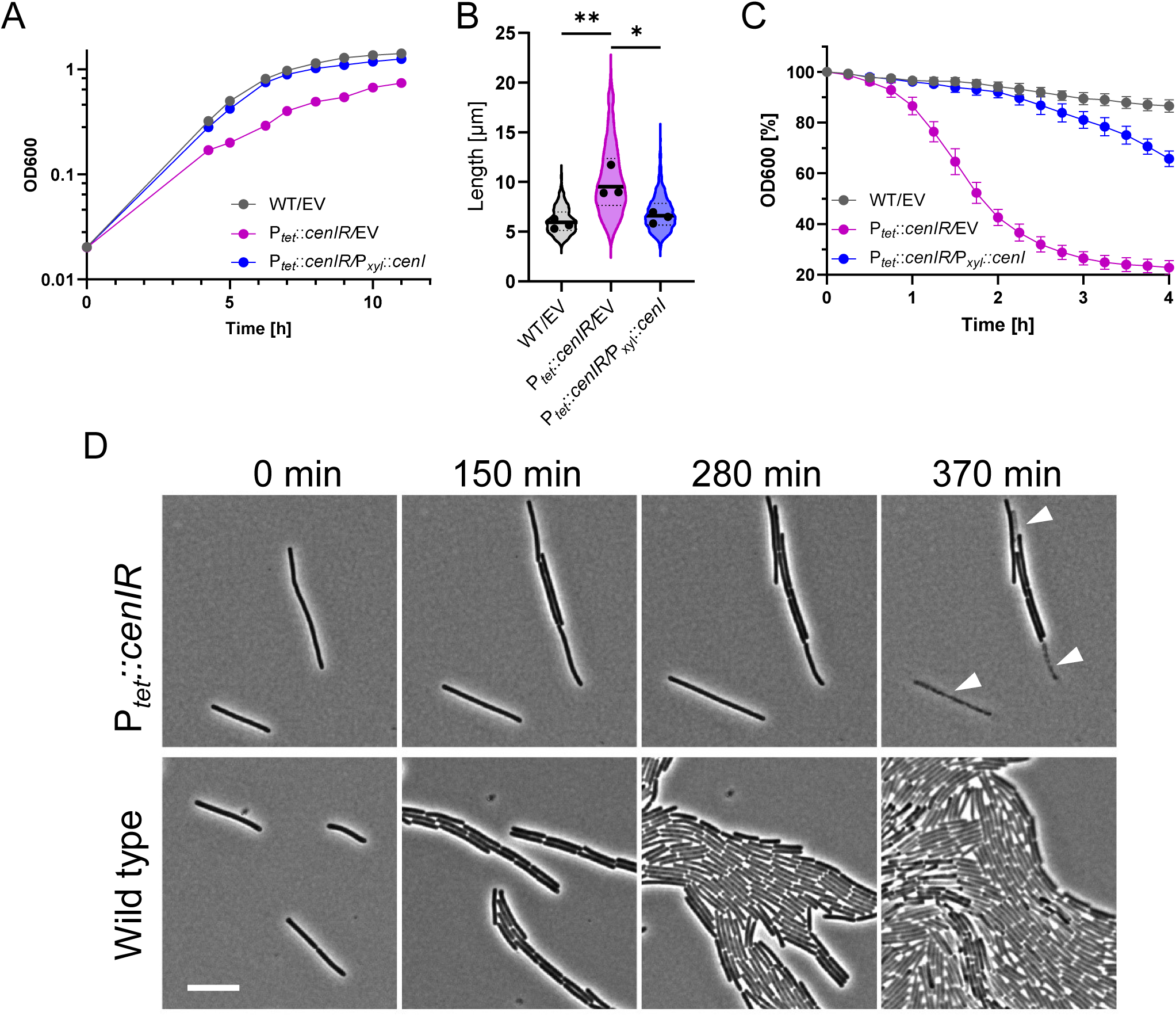
Depletion of *cenIR* causes slower growth, elongation and increased lysis. (A) Growth curves. Cultures that had been grown overnight in TY-thi with 2 ng/mL aTet were subcultured 1:50 into TY-thi with 1% xylose. At ∼5 h, cultures were sampled for phase contrast microscopy to determine cell length (B). At 6 h, the wildtype and the complemented strain were back-diluted 5-fold such that all strains would reach an OD_600_∼0.7 at 11 h, when cells were harvested for the lysis assay (C). Experiments were performed on three different days. Growth curves are representative. Cell lengths are graphed as violin plots using pooled data from all three experiments (≥200 cells per experiment). Bars and dotted lines are the mean and quartiles, respectively. Dots are the means of each experiment. Lysis is plotted as the mean ± SD from all three experiments. Statistical analysis was by one-way ANOVA (n = 3, * P ≤ 0.05, ** P ≤ 0.01). (D) Select images from time-lapse microscopy of the CenIR depletion strain and wild-type, both grown in the absence of aTet. Note that the depletion strain grew slower, elongated and eventually lysed (white carets). Size bar: 10 µm. Strains: WT/EV (UM1173), P*_tet_*::*cenIR/*P*_tet_::cenIR* (MK212), P*_tet_*::*cenIR*/EV (MK230), P*_tet_::cenIR* (UM1364).

**FIG 3.**
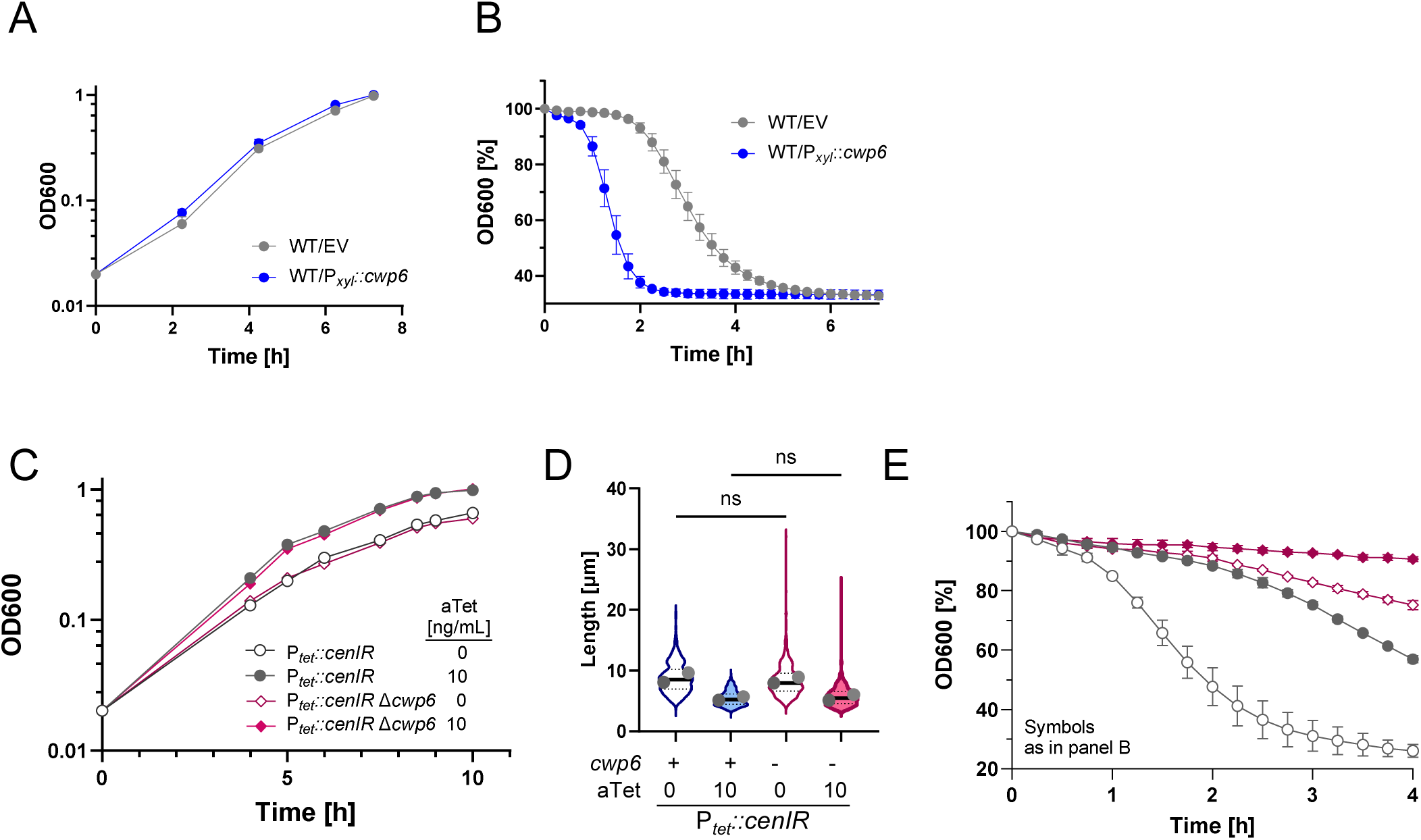
Cwp6 affects lysis but not growth rate or cell length. (A, B) Effect of *cwp6* induction on growth and lysis. Wildtype harboring empty vector (EV) or a P*_xyl_*::*cwp6* expression plasmid were grown in TY-thi with 1% xylose. At 7 h samples were harvested for the lysis assay. The growth curves are representative of three experiments. Lysis is graphed as the mean ± SD of three experiments. (C-E) Effect of deleting *cwp6* on growth, cell length and lysis. Strains grown overnight in TY with 2 ng/mL aTet were subcultured 1:50 into TY with 0 or 10 ng/mL aTet. Samples were harvested at 6.5 h for phase contrast microscopy to determine cell length. At 6 h, the 10 ng/mL aTet culture was back diluted ∼5-fold so that all strains had an OD_600_ = ∼0.7 at 10 h, when cells were harvested for the lysis assay. The growth curves are representative of two experiments. Cell lengths are graphed as violin plots with data pooled from two experiments (≥200 cells each). Bars and dotted lines show the means and quartiles, respectively. Circles show the means of each experiment. The indicated differences are not statistically significant (one way ANOVA, n = 2). Lysis is graphed as the mean ± SD of the two experiments. Strains: WT/EV (UM1173), WT/P*_xyl_::cwp6* (UM1352), P*_tet_*::*cenIR* (UM1364), P*_tet_*::*cenIR* Δ*cwp6* (MK237).

The phenotypic consequences of CenIR-depletion were confirmed using CRISPR interference (CRISPRi, (7)) against *cenIR*. Although viability in a spot titer assay was unaffected, cells scraped from the plates were elongated and showed signs of lysis (Fig. S2B).

Viability in the spot titer assay is difficult to reconcile with our inability to delete *cenIR*, although residual *cenIR* expression under depletion conditions might ameliorate phenotypic defects. Importantly, CenIR-depletion caused lysis in multiple assay formats (lysis assay, spot titer, time-lapse microscopy), and as will be shown below, we were eventually able to delete *cenIR* once we identified and removed the genes responsible for lysis. Thus, we consider *cenIR* to be essential or quasi-essential. A similar discrepancy emerged from large-scale Tn mutagenesis, where cenIR was classified as essential in an early study (5) but not in a more recent one (8).

### The CenIR regulon includes cell envelope genes

We speculated that identifying the genes in the CenIR regulon would shed light on its physiological function. To this end, we determined transcriptional profiles with and without CRISPRi knockdown of *<u>cenIR</u>*. CRISPRi was chosen over the P*_tet_*-based depletion strain because pilot studies showed that the high aTet required to phenocopy WT inhibits growth. Note that we use the term regulon loosely here and include genes that respond directly or indirectly to CRISPRi against *cenIR*.

Strains of *C. difficile* R20291 harboring either a CRISPRi plasmid targeting *cenI* or a negative control plasmid that expresses a non-targeting sgRNA were grown under CRISPRi-inducing conditions (i.e., 1% xylose). Cultures were maintained by repeated back-dilution for roughly 12 doublings, long enough to see a mild effect on growth rate (Fig. S2C). Cells were processed for RNA-seq as described previously (9).

RNA-seq analysis identified 14 genes whose transcript abundance changed by ≥4-fold (all p-values were ≤ 10^-8^) upon CRISPRi silencing of *cenIR* (Tables 1, S1). Only two of these genes were repressed, cenI and cenR, which were down 4- and 7-fold, respectively. Repression of these genes demonstrates that CRISPRi worked as intended and was specific for cenI (which is polar onto *cenR*) without any strong off-target effects. Consistent with CenI being a repressor, all other affected genes exhibited increased transcript levels, mostly in the range of 4- to 8-fold. A remarkable exception is cdr_0474, which is located 215 bp downstream of *cenIR* (Fig. 1) and for which mRNA abundance increased by 500-fold. CDR_0474 is a small protein of 136 amino acids found exclusively in *C. difficile*. It is predicted to have an N-terminal signal sequence, implying it is exported. Consistent with this, CDR_0474 has been detected in the secreted proteome (10). BLAST searches failed to identify similarity to any other proteins.

Genes that were derepressed at more moderate levels fall into three categories - regulation, metabolism and peptidoglycan (PG) modification. (i) SinR and SinR’ are regulatory proteins that in *C. difficile* have pleiotropic effects on sporulation, toxin production, and motility (11). (ii) Genes *cdr_2227* through *cdr_2241* are part of a large gene cluster involved in fermentation of succinate and glycine. All additional genes in this cluster were also induced, although by less than the 4-fold cutoff for inclusion in Table 1 (Table S1). (iii) Lastly, *cdr_2672* codes for Cwp6, a PG hydrolase (N-acetylmuramoyl-L-alanine amidase) and cdr_2797 codes for Ldt1, a L,D-transpeptidase that makes 3-3 crosslinks in PG. In view of the morphological defects we observed upon CenIR-depletion, we chose to follow-up on the cell envelope-associated genes *cdr_0474*, cwp6 and ldt1.

**TABLE 1.** Genes with fourfold change or more in transcript levels upon CRISPRi silencing *cenIR*.

| **Locus** | **Gene** | **Annotation** | **Log<sub>2</sub> fold change** | **P adjusted** |
| --- | --- | --- | --- | --- |
| *cdr_0472* | *cenI* | *blaI*-like | -2.06 | 2.7E-08 |
| *cdr_0473* | *cenR* | *blaR*-like | -2.82 | 2.0E-21 |
| *cdr_0474* |  | putative exported protein | 9.11 | 0.0E+00 |
| *cdr_2121* | *sinR* | HTH-type transcriptional regulator | 2.99 | 3.5E-20 |
| *cdr_2122* | *sinR'* | HTH-type transcriptional regulator | 2.72 | 6.6E-18 |
| *cdr_2227* | *macA* | 4-hydroxybutyrate dehydrogenase | 2.15 | 3.6E-11 |
| *cdr_2230* | *abfD* | gamma-aminobutyrate metabolism dehydratase/isomerase | 2.04 | 6.7E-09 |
| *cdr_2231* | *sucD* | succinate-semialdehyde dehydrogenase | 2.06 | 1.5E-08 |
| *cdr_2232* | *cat1* | succinyl-CoA:coenzyme A transferase | 2.16 | 1.2E-09 |
| *cdr_2233* | *yidE* | putative succinate transporter | 2.29 | 6.5E-12 |
| *cdr_2238* | *grdC* | glycine reductase complex component C subunit beta | 2.01 | 2.4E-10 |
| *cdr_2241* | *grdE* | glycine reductase complex component B subunits alpha and beta | 2.09 | 7.9E-09 |
| *cdr_2672* | *cwp6* | N-acetylmuramoyl-L-alanine amidase | 2.90 | 3.6E-25 |
| *cdr_2797* | *ldt1* | L,D-transpeptidase 1 | 2.68 | 1.5E-22 |

**TABLE 2.** Strains used in this study.

| Strain | Genotype | Plasmid feature | Source |
| --- | --- | --- | --- |
| R20291 | Wild-type *C. difficile* strain from UK outbreak (ribotype 027) |  |  |
| CDE5470 | R20291 $P_{cdr_{0474}}::sluc^{opt}$ | | This study |
| KB852 | R20291 $\Delta cwp6$ | | This study |
| MK212 | R20291 $P_{tet}::cenIR/pMK49$ | $P_{xyl}::cenI$ | This study |
| MK216 | R20291 $P_{tet}::cenIR/pMK42$ | CRISPRi *cdr_0474-1* | This study |
| MK217 | R20291 $P_{tet}::cenIR/pMK43$ | CRISPRi *cwp6-1* | This study |
| MK218 | R20291 $P_{tet}::cenIR/pMK44$ | CRISPRi *cwp6-2* | This study |
| MK227 | R20291 $P_{tet}::cenIR/pIA68$ | CRISPRi *ldt1-1* | This study |
| MK228 | R20291 $P_{tet}::cenIR/pIA69$ | CRISPRi *ldt1-2* | This study |
| MK229 | R20291 $P_{tet}::cenIR/pIA102$ | CRISPRi *cdr_0474-2* | This study |
| MK230 | R20291 $P_{tet}::cenIR/pBZ101$ | $P_{xyl}$ negative control | This study |
| MK231 | R20291 $P_{tet}::cenIR/pIA34$ | CRISPRi negative control | This study |
| MK237 | R20291 $P_{tet}::cenIR \Delta cwp6$ | | This study |
| MK252 | R20291 $\Delta cdr_{0474}$ | | This study |
| MK254 | R20291 $P_{tet}::cenIR \Delta cdr_{0474}$ | | This study |
| MK260 | R20291 $P_{cdr_{0474}}::sluc^{opt}/pIA59$ | CRISPRi *cenI* | This study |
| MK261 | R20291 $P_{cdr_{0474}}::sluc^{opt}/pIA34$ | CRISPRi negative control | This study |
| MK330 | R20291 $\Delta cdr_{0474}/pIA34$ | CRISPRi negative control | This study |
| MK331 | R20291 $\Delta cdr_{0474}/pIA59$ | CRISPRi *cenI* | This study |
| MK333 | R20291 $\Delta cenIR \Delta cdr_{0474}$ | | This study |
| MK338 | R20291 $\Delta cdr_{0474} P_{cdr_{0474}}::sluc^{opt}/pIA59$ | CRISPRi *cenI* | This study |
| MK339 | R20291 $\Delta cdr_{0474} P_{cdr_{0474}}::sluc^{opt}/pIA34$ | CRISPRi negative control | This study |
| MK342 | R20291 $\Delta cwp6 \Delta cenIR$ | | This study |
| UM1173 | R20291/pBZ101 | $P_{xyl}$ negative control | This study |
| UM1352 | R20291/pIA141 | $P_{xyl}::cwp6$ | This study |
| UM1364 | R20291 $P_{tet}::cenIR$ | | This study |
| UM359 | R20291/pIA34 | CRISPRi negative control | (7) |
| UM469 | R20291/pIA59 | CRISPRi *cenI* | This study |

### Ldt1 is not linked to any of the morphological defects

We used CRISPRi to silence expression of *ldt1* in the P*_tet_*::*cenIR* depletion background. None of the morphological defects associated with CenIR depletion were affected (Fig. 4). We have shown elsewhere that one of the CRISPRi plasmids used in this experiment is effective at reducing Ldt1 levels, so the lack of phenotypic changes is not due to ineffective guides (12). This outcome is not too surprising because *C. difficile* has five *ldt* genes, of which three (including *ldt1*) show considerable functional redundancy and are alone sufficient for wild type growth rate and morphology (12). Based on these considerations, we decided not to pursue Ldt1 further by constructing a deletion mutant. We did, however, use anti-Ldt1 anti-serum to examine Ldt1 abundance by Western blotting. Depletion of CenIR increased Ldt1 levels about 2-fold (Fig. S3), less than the 4-fold increase indicated by RNA-seq but still in the same direction. In summary, Ldt1 is indeed elevated upon CenIR depletion, but this increase does not appear to be connected to any phenotypic changes.

**FIG 4.**
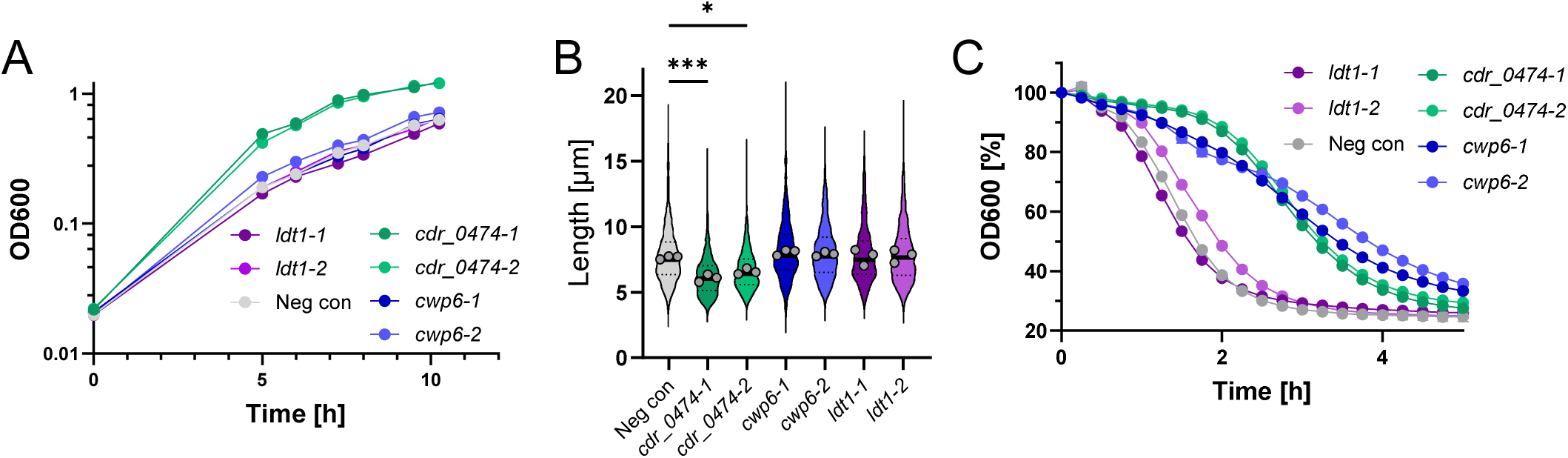
Effect of CRISPRi silencing of select regulon members on CenIR depletion phenotypes. (A) Growth curves. Strains grown overnight in TY-thi with 2 ng/mL aTet were subcultured 1:50 into TY-thi with 1% xylose and grown for about 10 h. At 8 h, cultures were sampled for microscopy to determine cell length (B). The two faster growing strains were back diluted 7-fold at 6 h to ensure that all strains were at an OD_600_∼0.65 at 10.5 h when cells were harvested for the lysis assay (C). Growth curves and lysis assay are representative of three experiments. Cell lengths are graphed as violin plots using pooled data from all three experiments (≥200 cells per experiment). The bar and dotted lines are the mean and quartiles. Dots are the mean of each experiment. Statistical analysis is by one-way ANOVA, n = 3 (* P ≤ 0.05, *** P ≤ 0.001). The strains shown are the P*_tet_*::*cenIR* depletion strain containing various CRISPRi plasmids with sgRNAs targeting the indicated gene. “Neg con” is a control with an sgRNA that does not target anywhere in the genome. The strains are: “Neg con” (MK231), *ldt1-1* (MK227), *ldt1-2* (MK228), *cdr_474-1* (MK216), *cdr_474-2* (MK229), *cwp6-1* (MK217), *cwp6-2* (MK218).

### Deletion of *cdr_0474* corrects phenotypic defects caused by loss of CenIR

Recall that *cdr_0474* is a gene of unknown function that according to RNA-seq was induced 500-fold upon CenIR depletion. To verify the large induction, we endeavored to build reporter plasmids by cloning the cdr_0474 promoter region upstream of codon-optimized genes for mCherry red fluorescent protein or NanoLuc luciferase (13–15). Despite repeated attempts, we were unable to recover intact plasmids, which might be toxic to *E. coli* due to depletion of rare tRNAs when the reporter genes are expressed at high levels. Ultimately, we were able to construct a P*cdr_0474*::*sluc^opt^* fusion in Tn916 in Bacillus subtilis. The reporter transposon was transferred by conjugation into C. difficile, where it integrated at the intergenic region upstream of *cdr_0029* as revealed by whole genome sequencing. CRISPRi silencing of *cenIR* induced the secreted NanoLuc reporter (14) about 100-fold (Fig. 6), confirming that the reporter strain responds as expected based on the RNAseq results. Induction of the reporter was not affected by the deletion of cdr_0474 (Fig. 6).

**FIG 5.**
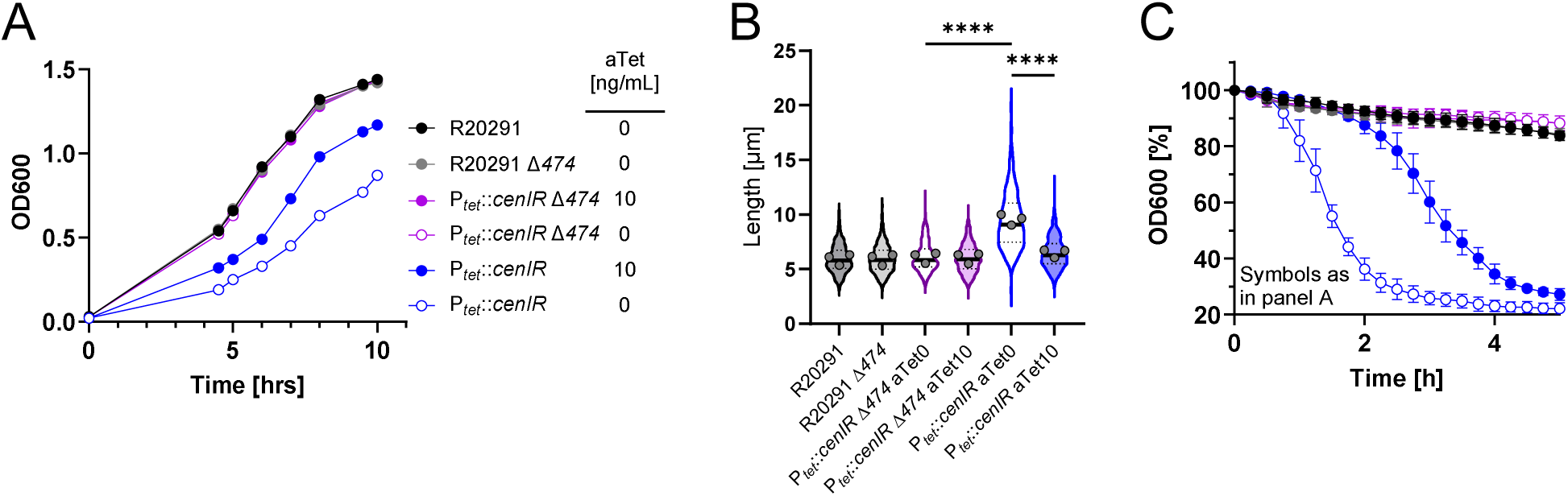
Deletion of *cdr_0474* rescues CenIR depletion phenotypes. A. Growth curves. Strains grown overnight in TY (wild type) or TY with 2 ng/mL aTet (P_tet_::*cenIR*) were subcultured 1:50 into TY (wild type) or TY ± 10 ng/mL aTet (P_tet_::*cenIR*) and growth was monitored for 11 h. At 7.5 h, cultures were sampled for microscopy to determine cell length (B). Faster growing strains were back diluted at 6 and 9.5 h to ensure that all strains were at an OD_600_∼0.65 when harvested for the lysis assay (C). Growth curves are representative of three experiments. Cell lengths are graphed as violin plots with pooled data from three experiments (≥200 cells each). Bars and dotted lines are the means and quartiles, respectively. Circles are the means of each experiment. Statistical analysis was by one-way ANOVA, n = 3 (**** P ≤ 0.0001). Lysis is graphed as the mean ± SD of three experiments. Strains: R20291, R20291 Δ*cdr_0474* (MK252), P*_tet_*::*cenIR* (UM1364), P*_tet_*::*cenIR* Δ*cdr_0474* (MK254).

**FIG 6.**
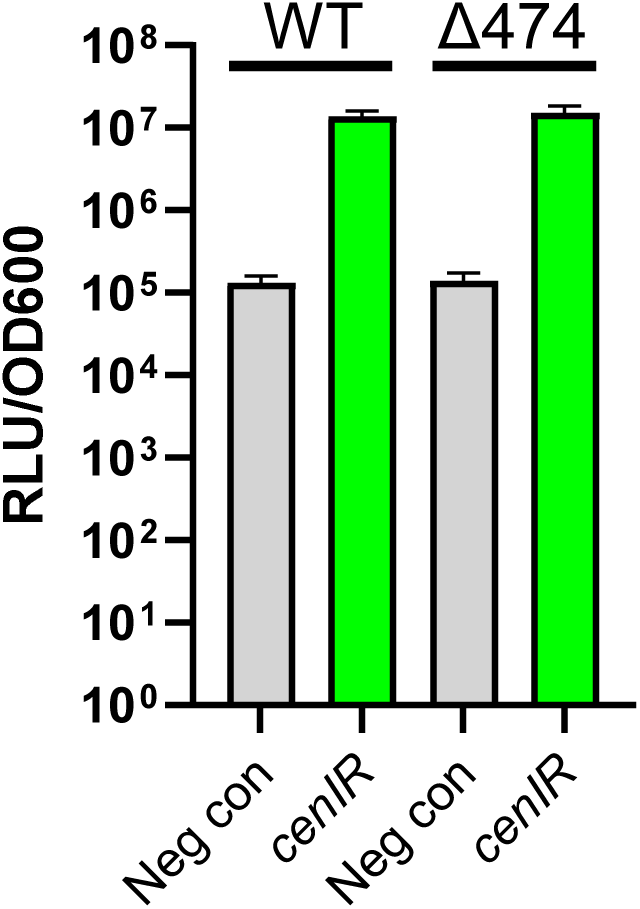
Depletion of CenIR induces P*_cdr_0474_*::*sluc^opt^*. The P*_cdr_0474_* reporter strain harboring a negative control plasmid or a CRISPRi plasmid against *cenI* was grown overnight in TY-Thi, subcultured 1:100 into TY-Thi with 1% xylose, grown to an OD_600_ of 0.5, and luciferase activity measured. When *cenI* was silenced, luciferase activity increased about 100-fold. Data are presented as the mean and SD from two experiments performed on different days with two technical replicates each. Strains: P*_cdr_0474_*-*sluc*^opt^/Neg con (MK261), P*_cdr_0474_*-*sluc*^opt^/sgRNA-*cenI* (MK260), Δ*cdr_0474* P*_cdr_0474_*-*sluc*^opt^/Neg con (MK338), Δ*cdr_0474* P*_cdr_0474_*-*sluc*^opt^/CRISPRi-*cenI* (MK339).

We next tested whether CDR_0474 contributes to any of the CenIR-depletion phenotypes. We started by introducing plasmids for CRISPRI knockdown of *cdr_0474* into the P*_tet_*::*cenIR* depletion background. Strikingly, knockdown of cdr_0474 by CRISPRi with either of 2 different sgRNAs improved growth rate, reduced cell length and slowed lysis as compared to a control CRISPRi plasmid with a non-targeting sgRNA (Fig. 4). The reversal of CenIR depletion phenotypes upon CRISPRi silencing *cdr_0474* led us to investigate the effect of deleting *cdr_0474*. Indeed, a *cdr_0474* deletion in the depletion background completely restored wild-type growth, cell length and lysis activity, while deletion of cdr_0474 in the wild-type background had no effect (Fig. 5). The mild increase of Ldt1 seen by Western blot upon CRISPRi silencing *cenIR* was also abolished in the Δcdr_0474 mutant (Fig. S3). Finally, we found that *cenIR* could be deleted in the Δcdr_0474 background. Both growth and lysis phenocopy the wild-type strain (Fig. S4A,B). Thus, the apparent lethality of deleting cenIR in an otherwise wild-type background is likely related to run-away expression of cdr_0474.

### Cwp6 is responsible for the lysis phenotype

Elevated production of Cwp6, an annotated PG hydrolase, seemed likely to underlie the propensity for lysis when C. difficile is depleted of CenIR. To test this hypothesis, we overexpressed cwp6 from a P_xyl_ plasmid in the wild-type background. Although growth rate was not affected, cells lysed markedly faster in the Triton X-100 based lysis assay (Fig. 3A). Conversely, deleting cwp6 in the P*_tet_*::*cenIR* depletion background ameliorated the lysis phenotype without affecting growth or cell length (Fig. 3B-D). Similar results were obtained using CRISPRi to knock down expression of cwp6 in the P*_tet_*::*cenIR* depletion background (Fig. 4). Based on these results, we attempted to delete *cenIR* in the Δ*cwp6* background using the same sgRNA and homology repair regions that failed to generate a *cenIR* deletion in the wild-type background. Deletions were easily obtained. The resulting Δ*cenIR* Δ*cwp6* mutant grew like wild-type, with no cell elongation or evidence of lysis (Fig. S4C,D). These results are consistent with Cwp6 causing the increased lysis seen upon depletion of *cenIR*. Moreover, the essentiality of cenIR is due to the lysis phenotype, not slower growth or elongation.

### Unsuccessful efforts to identify conditions that induce the *cenIR* regulon

We used our chromosomal P*_cdr_0474_*::*sluc^opt^* fusion to screen for conditions or signals that activate the regulon. Antibiotics were tested by growing the reporter strain in TY broth with a range of antibiotic concentrations. The antibiotics tested included two β-lactams (cefalexin, cefoperazone), additional cell wall targeting drugs (bacitracin, ramoplanin, ristocetin) and a DNA gyrase inhibitor (novobiocin) strain. A strain harboring the CRISPRi-cenI plasmid served as a positive control and showed 100-fold induction of luciferase activity in the presence of 1% xylose. None of the antibiotics affected luciferase even 1.5-fold. The lack of response to β-lactams is not surprising since CenR lacks a β-lactam-binding domain at the C-terminus, but we still considered it possible that CenIR might respond to cell envelope damage induced by β-lactams or other cell wall targeting drugs. Either this is not the case, or we have yet to test the right compounds. Extending our speculation that the CenIR system might respond to perturbations in the cell wall, we assayed P_cdr_0474_::sluc^opt^ expression after CRISPRi silencing a dozen cell envelope-associated genes (pbp1, pbp2, fabK, murJ2, zapA, ddl, ispG, dxr, pgm2, murB, slpA, and walA) (8). Strains harboring the respective CRISPRi plasmids were subcultured into TY-Thi with 1% xylose and sampled after 5.5h. The positive control, CRISPRi-cenI induced the reporter about 100-fold, while none of the other constructs increased luciferase activity even 1.5-fold.

### Deletion of *cenIR* does not affect the MIC of select antibiotics

Once it became possible to delete cenIR (by first deleting cdr_0474 or cwp6), we could ask whether this regulator is important for responding to antibiotics. To this end, we used minimum inhibitory concentration (MIC) assays to compare wild type R20291, Δcdr_0474 and the Δcdr_0474 ΔcenIR double mutant. We tested three β-lactam antibiotics (ampicillin, methicillin, meropenem), two cell wall-acting antibiotics from different classes (bacitracin, vancomycin) and a replication inhibitor (novobiocin). However, sensitivities to these antibiotics were the same for all three strains (Table S2).

### BlaR homologs without a predicted transpeptidase domain are widespread

At the outset of this study we thought the absence of a β-lactam-binding TP domain in CenR made this sensor an unusual BlaR-like protein. Close inspection of the six other BlaR-like proteins in strain R20291 revealed that while all are predicted membrane proteins with an M56 peptidase domain (PF05569) only one of them has a TP domain (PF00905) (Table S3). That one is the previously characterized canonical BlaR protein CDR_0412 (4). The remaining six BlaR-like proteins, including CenR, have unique C-terminal domains that are much shorter than a TP domain and in most cases predicted to be largely unstructured (Table S3). Two of these novel BlaR-like proteins are encoded in an operon with a corresponding blaI-like encoding gene (cdr_2276/2277 and cdr_2279/2280). Interestingly, the latter pair is upstream of CDR_2278, which has a PG binding domain typically found in PG hydrolases (PF01471), suggesting that it might play a role in regulating cell wall homeostasis.

These observations made us wonder whether it might be common for a BlaR-like protein to sense something other than a β-lactam antibiotic. We identified one example, a non-canonical BlaR homolog (i.e., without the β-lactam-binding TP domain) that has been studied in both Mycobacterium tuberculosis and in Mycobacterium abscessus (16, 17). In M. abscessus it has been linked to energy metabolism rather than antibacterial resistance. To identify additional atypical BlaR proteins, we used the InterPro database to retrieve roughly 15,000 protein entries with an M56 domain (PF05569), then grouped these by domain architecture (18, 19). There were 254 different domain arrangements of which 9 included 100 or more entries (Fig. 7). The most abundant architecture is for an M56 domain followed by C-terminal regions of various lengths but with no identifiable domain. The second most common architecture has a C-terminal TonB_C domain. Bona fide BlaR proteins with a TP domain that can bind β-lactams came in third on the list and represented only ∼6% of “BlaR-like” homologs.

**FIG 7.**
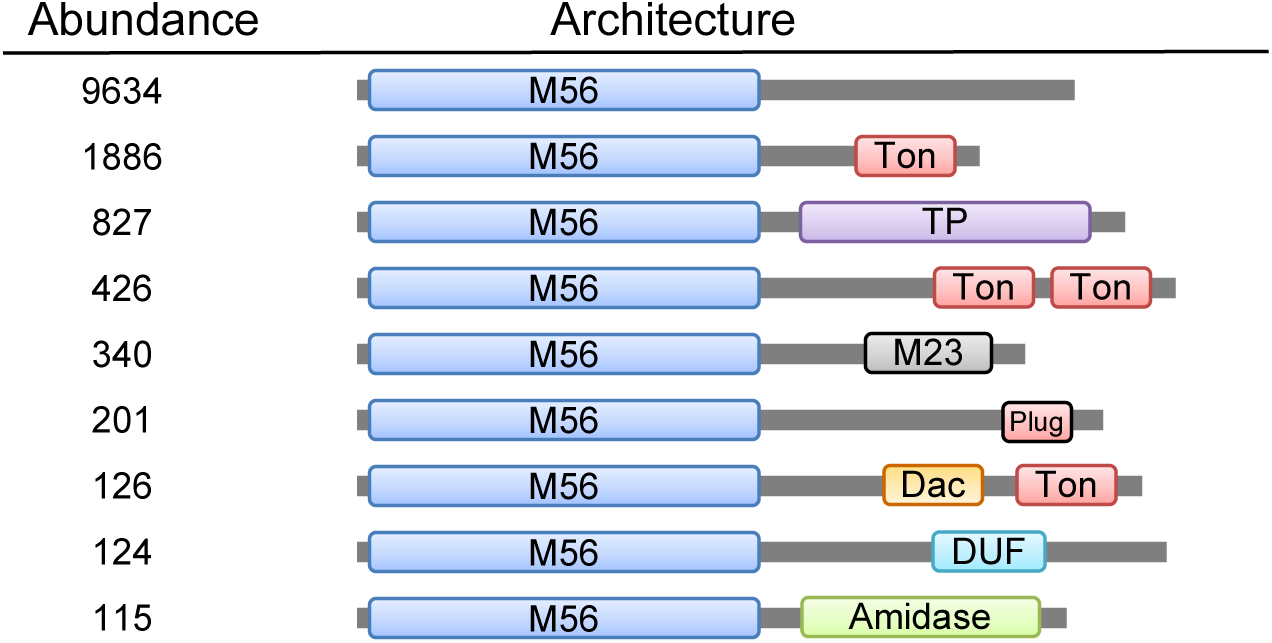
M56 domain proteins have diverse C-terminal domains. The M56 domain (PF05569) was used to search the InterPro database for homologs. The roughly 15,000 hits grouped into 254 domain architectures. Architecture arrangements with 100 or more entries are shown. Abbreviations are as follows. Ton: TonB_C domain (PF03544), TP: transpeptidase domain (PF00905), M23: Peptidase M23 domain (PF01551), Plug: TonB dependent receptor plug domain (PF07715), Dac: CarbopepD_reg_2 domain (PF13715), DUF: Domain of unknown function 4825 (PF16107), Amidase: Amidase_3 domain (PF01520).

To verify that the BlaR homologs included in this analysis are sensors involved in regulating gene expression, and not, for example, proteins that have an M56 peptidase domain for some other reason, we looked for a closely linked BlaI-like protein. This analysis was performed using the EFI - Genome Neighborhood Tool (20, 21). About 96% of the BlaR homologs were within 5 genes of a predicted BlaI-like transcriptional repressor. In most cases, the BlaI homolog was upstream of the BlaR homolog.

## Discussion

Here we have identified and characterized a BlaIR-like regulatory system composed of the predicted transcriptional repressor CenI and the predicted membrane metalloprotease CenR. Unlike canonical BlaIR-like systems, which sense β-lactams to induce expression of a β-lactamase, CenIR is not implicated in β-lactam resistance. This conclusion is supported by three observations. (i) Deletion of CenIR had no effect on β-lactam resistance. (ii) The CenIR regulon does not include any predicted β-lactamases. (iii) The extracellular sensing domain of CenR is not a penicillin-binding domain. Of note, C. difficile has a conventional BlaIR pair that mediates induction of a β-lactamase (4); this BlaIR pair is distinct from CenIR.

CenIR caught our attention because a large-scale transposon mutagenesis indicated it is essential for vegetative growth (5). Essentiality is interesting because it is unexpected. BlaIR systems typically mediate β-lactam resistance, so they are dispensable under standard growth conditions. Indeed, CenIR scored as non-essential in our own Tn-seq analysis of C. difficile (8). Nevertheless, we were unable to delete the locus. To gain insight into the importance and role of CenIR, we used two depletion strains, one based on CRISPRi, the other on replacing the native promoter with P_tet_. In both cases, depleting cells of CenIR resulted in slower growth, elongation and eventual cell lysis.

RNA-seq suggests the CenIR regulon is small, about 12 genes. All of these genes were induced upon CenIR depletion, as expected if CenI is a repressor. Because the experimental design involved targeting cenIR with CRISPRi, it is not possible to say whether cenIR expression is autoregulated. The most interesting gene in the regulon is cdr_0474, which is located 215 bp downstream of cenIR and was induced up to 500-fold. CDR_0474 is a small exported protein (10) unique to C. difficile. Its function is unknown, and its sequence offers no clues as to what it might be doing. Baseline levels of sequence reads in RNA-seq for CDR_0474 indicate expression is high even before depletion of CenIR (Table S1). High basal expression suggests CDR_0474 plays a structural role rather than a catalytic one.

cdr_0474 is not essential by transposon sequencing and we had no difficulty deleting the gene. Importantly, deleting cdr_0474 suppressed the morphological and lysis defects associated with CenIR depletion, and it was now possible to delete cenIR. The CenIR regulon also includes two enzymes involved in peptidoglycan biogenesis, ldt1 and cwp6, which were upregulated in RNA-seq by 6- and 7-fold, respectively. Ldt1 is a peptidoglycan crosslinking enzyme. Follow-up experiments with Ldt1 failed to link it to any morphological defects. Cwp6 is a peptidoglycan hydrolase. Follow-up experiments revealed Cwp6 mediates the lysis phenotype, but not slower growth or cell elongation. Importantly, deleting cwp6 rendered CenIR no longer essential.

Based on these findings, we propose a model for essentiality of CenIR. In this model, CenI represses cdr_0474, which is highly induced in the absence of CenI. Excess CDR_0474 protein disrupts normal envelope biogenesis, which leads to increased expression of the cell wall amidase cwp6. Lethality is due to Cwp6-mediated lysis. We attempted to test this hypothesis by overexpressing cdr_0474 from a plasmid. However, we did not find any viability or morphology defects, likely because we were unable to achieve high enough levels of expression.

The physiological role of CenIR remains an open question. The location of CenR in the membrane, the morphological defects observed upon CenIR depletion and the genes in the CenIR regulon all point to a role in cell envelope biogenesis or homeostasis. Nevertheless, we were unable to identify any antibiotics or mutations that induced a P_cdr_0474_::sluc^opt^ reporter, nor did deletion of CenIR in the *Δcdr_0474 background* sensitize *C. difficile* to any of the antibiotics we tested. By analogy to canonical BlaIR pairs, whatever signal activates CenR does so by binding to its C-terminal tail. Prediction programs indicate this region (amino acids 334-460) is mostly unstructured, although a small globular domain comprising amino acids 417-459 is weakly predicted by AF3 (Fig. S5). Unstructured domains can serve as tethers, binding sites for proteins, and probes for gaps in the peptidoglycan wall (22–26).

Interestingly, bioinformatic analysis of M56 domain proteins argues that most “BlaIR-like” regulatory systems are not likely to be involved in β-lactam resistance. Only one of *C. difficile’s* seven annotated BlaIR-like gene pairs has a BlaR protein with a C-terminal penicillin-binding domain (4). The remaining six BlaR proteins have largely unstructured C-terminal domains that are not homologous to one another. C. difficile is not an outlier in this regard. Of the roughly 15,000 BlaIR systems identified in the InterPro database, only about 6% are predicted to bind a β-lactam. Putative signal-binding domains in the remaining BlaR-like proteins are for the most part diverse and it is not possible to predict what ligands they bind. We infer that BlaIR-like systems couple many different signals to changes in gene expression via a conserved mechanism involving regulated proteolysis of a transcriptional repressor protein.

## Materials and Methods

### Strains, media, and growth conditions

Bacterial strains used in this study are listed in Table 2. *C. difficile* strains were derived from R20291 (27). C. difficile was routinely grown in tryptone-yeast extract (TY) medium, supplemented as needed with thiamphenicol at 10 μg/ml (TY-Thi). TY medium consisted of 3% tryptone, 2% yeast extract, and 2% agar (for plates). Brian heart infusion (BHI) media was prepared per manufacturer’s (DIFCO) instructions. BHIS media is BHI supplemented with 5 g/L yeast extract and 4 g/L glucose. C. difficile strains were maintained at 37°C in an anaerobic chamber (Coy Laboratory Products) in an atmosphere of 2% H_2_, 5% CO_2_, and 93% N_2_. Escherichia coli strains were grown in LB medium at 37°C with chloramphenicol at 10 μg/ml and/or ampicillin at 100 μg/ml as needed. LB medium contained 1% tryptone, 0.5% yeast extract, 0.5% NaCl, and 1.5% agar (for plates). OD_600_ measurements were made with the WPA Biowave CO8000 tube reader in the anaerobic chamber.

### Plasmid and strain construction

Plasmids are listed in Table S4 and were constructed with HiFi DNA Assembly from New England Biolabs (Ipswich, MA). Oligonucleotide primers (Table S5) were synthesized by Integrated DNA Technologies (Coralville, IA). Regions that were constructed by PCR were verified by DNA sequencing. Plasmids were propagated in E. coli HB101/pRK24 and conjugated into C. difficile R20291 according to (9). Details relevant to plasmid construction are provided in Table S4.

### Strain construction by CRISPR editing

Gene deletions were achieved by CRISPR editing, using pIA123 as the parental vector (Addgene #234038). This plasmid carries a xylose-inducible cas9 gene, an easily swapable sgRNA (MscI-NotI sites) under control of a constitutive promoter, as well as a unique PstI site for insertion of the homology repair regions (typically 500 bp upstream and 500 bp downstream of the desired deletion). We generally added the homology region first, followed by insertion of the sgRNA. We typically screened 3-5 sgRNA candidates for optimal killing upon induction of cas9. Constructs with different guides were conjugated into the appropriate C. difficile strain and grown overnight in TY-Thi. The overnight culture was serially diluted (10-fold dilutions), spotted on both TY-Thi and TY-Thi 1% xylose plates and growth evaluated after 17h incubation at 37°C. A suitable guide will achieve about 10^4^-fold loss of viability upon plating on xylose. Strains harboring an editing plasmid with an optimal sgRNA were then plated for selection by spreading 100 µL of a diluted overnight culture (10^-1^ and 10^-2^ dilution) on TY-Thi 1% xylose. After overnight growth, ∼8 colonies were restruck on TY 1% xylose. The next day, colony PCR was used to evaluate which candidates have the intended deletion. Successful candidates were then restruck for multiple rounds on TY to achieve plasmid loss, as evaluated by patching on TY versus TY-Thi plates. CRISPR editing was also used to insert the P_tet_ promoter elements upstream of the pepIR operon. In that case the sequence for tetR and regulatory elements was included between the homology regions.

### Colony PCR

To prepare template DNA for PCR, a single colony is picked into 50 µL ThermoPol reaction buffer (NEB) with 0.5 µL thermolabile ProteinaseK (NEB), incubated at 37°C for 30 min, then the proteinase is heat deactivated for 10 min at 55°C. Of this, 1 µL is used as template in subsequent PCR.

### Construction of a P*_cdr_0474_-sluc^opt^*into reporter strain using a modified Tn916 transposon

Because the cdr0474 promoter was toxic in E. coli, a reporter fusion was constructed in a Tn916 transposon directly in Bacillus subtilis BS49 (28). The reporter construct placed 300 nucleotides upstream of cdr_0474 in front of the gene for secreted NanoLuc, sluc^opt^ (14). Five DNA fragments were amplified by PCR using the primers listed in Table S5. These fragments were: 1) Tn916 Frag1 (primers 7889/7890; template pSMB47 (29)), 2) the P_cdr0474_ promoter region (primers 7891/7892, template R20291 genomic DNA), 3) the sluc^opt^ reporter gene (primers 7893/7894; template pAP24 (14)), 4) a spectinomycin resistance cassette (primers 7895/7896; template pDG1730 (30)), and 5) Tn916 Frag2 (primers 7897/7898; template pSMB47). The purified PCR products were combined by isothermal assembly and the resulting construct was directly transformed into competent B. subtilis BS49. Transformants were selected on agar medium containing spectinomycin at 100 µg/ml. To confirm correct assembly and insertion of the reporter cassette into the Tn916 element, it was amplified by colony PCR and the PCR products were analyzed by Sanger sequencing. A confirmed Tn916 reporter construct was transferred from B. subtilis to C. difficile R20291 by conjugations as previously described (31). Transconjugants were selected on BHIS plates containing tetracycline (10 µg/ml), cefoxitin (8 µg/ml), and kanamycin (50 µg/ml). The presence of the P_cdr_0474_-sluc^opt^ reporter in the final strain, CDE5470, was confirmed by PCR. Whole genome sequencing of the final reporter strain determined the insertion to be in the intergenic region between acoL and cdr20291_0029 at position 63422-63425.

### Microscopy

Cells were immobilized using thin agarose pads (1%) as described (15). Phase-contrast micrographs were recorded on an Olympus BX60 microscope equipped with a 100× UPlanXApo objective (numerical aperture, 1.45). Micrographs were captured with a Hamamatsu Orca Flash 4.0 V2+ complementary metal oxide semiconductor (CMOS) camera. The image analysis tool MicrobeJ (32) was used to measure cell length. A minimum of 200 cells were measured for each condition.

### Live cell microscopy

Growing cells were imaged anaerobically according to (33) with the following modifications. Gene frames (size 125 µL from Thermo Scientific) attached to a microscope slide, 2% molten agarose (RPI) and 2X TY were brought into the anaerobic chamber the day before use so they had time to become anaerobic. The agarose was kept at 68°C. Shortly before assembling the slides, the 2X TY was also warmed to 68°C, then mixed 1:1 with the molten agarose, and FM4-64 was added to a final concentration of 1 µg/mL. 500 µL of this mixture was pipetted into the Gene frame well and covered with a second microscope slide. After about 5 min for the agarose to solidify, the microscope slide cover was removed and bacterial samples were spotted on to the agarose pad. The slide was then sealed with a 24 x 50 mm cover slip and removed from the chamber. The samples were kept at 37°C and imaged every 5 min by phase contrast and red fluorescence microscopy with an inverted Nikon Ti2 Eclipse as described (34).

### Lysis assay

Lysis activity was determined as reported (9). Briefly, cell cultures (1 mL) were removed from the anaerobic chamber, pelleted, and resuspended in 700 mL of 0.01% Triton X-100 in 50 mM NaPO4 buffer (pH 7.4). Of this, three 200-mL replicates were pipetted into wells of a clear, flat-bottom 96-well plate. The turbidity was measured at 600 nm every 15 min for 10 h at room temperature in a plate reader (Tecan Infinite M200 Pro). Note that lysis activity is growth phase dependent (Fig. S1), so cultures were sampled for the assay at the same OD_600_, typically 0.6-0.8.

### Western blot

Expression level of Ldt1 was quantitated by Western blotting as described (12). Details for culture growth are given in the legend of Fig. S2.

### Culture growth and RNA isolation for RNAseq

Two independent cultures each of R20291/pIA59 (CRISPRi-cenIR) and of the negative control strain R20291/pIA34 were grown overnight in TY-Thi 1% NaCl. The next day, these were subcultured 1:30 into TY-Thi 1% NaCl containing 1% xylose and grown for 13h. Cultures were kept in late log phase by monitoring the OD_600_ and twice back-diluting 5-fold once they had reached an OD_600_ of 0.5 (Fig S2). Cells were fixed by rapidly mixing 25 mL culture with 25 mL ice-cold 1:1 ethanol-acetone that had been brought into the anaerobic chamber on dry ice. Fixed cells were stored at -80°C until further workup. RNA was isolated according to (9).

### Transcript profiling (RNAseq)

RNA samples were submitted to the Microbial Genome Sequencing Center (MiGS) in Pittsburgh, PA. MiGS performed Illumina stranded RNA library preparation paired with RiboZero Plus (per the manufacturer’s specifications) and sequenced on a NextSeq 500 using a 75cyc high-output flow cell. Fastq files were trimmed and filtered using a combination of Trimmomatic (35) and FastQC (36). Alignment, normalization, and differential expression were analyzed with LaserGene v18.1.0.426. Annotation was imported from NCBI with additional information obtained from proGenomes (37). All subsequent analysis was performed in Excel and GraphPad Prism 9.

### Luciferase assays

Luciferase activity was measured with Nano-Glo reagents from Promega (Madison, WI). Working stock of the assay reagent refers to a 1:50 dilution of the Nano-Glo substrate in assay buffer. Immediately before use, the working stock was further diluted 1:100 in water. Bacterial cultures (50 or 100 µL) were added to a white 96-well plate, mixed with an equal volume of the diluted Nano-Glo substrate, and allowed to sit 5 min in the dark at room temperature. Luminescence was measured in a plate reader (Tecan Infinite M200 Pro), and reported as relative light units (RLU) normalized to cell density.

### Minimum inhibitory concentration

Susceptibility to antibiotics was determined in biological duplicate on two separate days as described (9).

### Bioinformatics

The InterPro database (Feb 2026) was searched for entries with an M56 domain (PF05569), then grouped by domain architecture (18, 19). To determine whether BlaR homologs were found adjacent to BlaI homologs, the roughly 15,000 hits from InterPro were downloaded and analyzed with bioinformatics tools at EFI (Enzyme Function Initiative, version Feb 2026) (20, 21). First a sequence similarity network (SSN) is created with the Enzyme Similarity Tool (EFI-EST) using the default settings and finalizing without alignment threshold or length restriction. The SSN is transferred to the EFI Gene Neighborhood Tool (GNT) with the Neighborhood size set to within 5 genes. The resulting PFAM family co-occurrence table was used to calculate the frequency of finding PF05569 within 5 genes of BlaI (PF03965).

## Data availability

RNA-seq data were submitted to the NCBI GEO repository and assigned accession number GSE327362.

## Acknowledgments

This work was supported by Public Health Service Grant R01 AI155492 (C.D.E. and D.S.W). We thank Kevin Bollinger and Paige Brannen for strain KB852 and plasmid pPB103. We thank Jeroen Corver and Paul Hensbergen from the Leiden University Medical Center, and members of the Ellermeier and Weiss laboratories for helpful discussions.

